# Microfluidic platform for live cell imaging of 3D cultures with clone retrieval

**DOI:** 10.1101/2020.02.17.952689

**Authors:** Carla Mulas, Andrew C Hodgson, Timo N Kohler, Chibeza C Agley, Florian Hollfelder, Austin Smith, Kevin Chalut

**Affiliations:** Wellcome-MRC Cambridge Stem Cell Institute, Jeffrey Cheah Biomedical Centre, University of Cambridge, Puddicombe Way, Cambridge Biomedical Campus, Cambridge, CB2 0AW, UK; Department of Biochemistry, University of Cambridge, 80 Tennis Court Road, Cambridge, CB2 1GA, UK; Living Systems Institute, University of Exeter, Exeter, EX4 4QD, UK; Department of Physics, University of Cambridge, 19 J J Thomson Avenue, Cambridge, CB3 0HE, UK

## Abstract

Combining live imaging with the ability to retrieve individual cells of interest remains a technical challenge. These combined methods are of particular interest when studying highly dynamic or transient, asynchronous or heterogeneous cell biological and developmental processes. Here we present a method to encapsulate live cells in a 3D hydrogel matrix, via droplet compartmentalisation. Using a small-scale screen, we optimised matrix conditions for the culture and multilineage differentiation of mouse embryonic stem (ES) cells. Moreover, we designed a custom microfluidic platform that is compatible with live imaging. With this platform we are able to retain or extract individual bead/droplets by media flow only, obviating the need for enzymatic cell removal from the platform. We show that we can differentiate mES cells, monitor reporter expression by live imaging, and retrieve individual droplets for functional assays, correlating reporter expression with functional response. Overall, we present a highly flexible 3D cell encapsulation and microfluidic platform that enables both monitoring of cellular dynamics and retrieval for molecular and functional assays.

## INTRODUCTION

Asynchrony and heterogeneity are key challenges when studying dynamic processes in biology, such as changes in cell state. Heterogeneity can be dissected by following individual cells over time using live imaging of reporter systems. However, such studies are limited by the number of parameters that can be observed at the same time. We can significantly increase the number of parameters, and more accurately infer cellular state, by using single cell RNA sequencing. Yet, in transcriptomic analysis, the history of a cell is typically lost and we can only place cells in ‘pseudo-time’ by making significant assumptions about the trajectories cells follow. On the other hand, microfluidic platforms have emerged which promise to overcome the limitations mentioned above and integrate live imaging with mRNA sequencing (for example^1,2^). These systems, however, often lack versatility, so that it is not possible to recover cells for functional assays, such as testing differentiation potential. Furthermore, given the need for most cells to grow in adherent conditions, enzymatic methods, which require inactivation steps, are often used to extract cells from the device.

Combining live imaging with cell retrieval for functional and molecular analysis could greatly help us better understand dynamic cellular process in controlled conditions. We have designed a highly flexible cell encapsulation system and microfluidic platform, and validated it using ES cell differentiation as a model system. ES cells possess the remarkable ability to divide indefinitely while maintaining the capacity to give rise to all cell types of the adult organism. Exit from the ES cell state and initiation of differentiation is a poorly understood key step. The study of this transition is hindered by a large degree of temporal heterogeneity ^3,4^. Indeed, at any given point in time during exit from the ES cell state, cells can be in different stages of the differentiation process. This asynchrony has confounded our understanding of how ES cells change state and initiate multilineage differentiation.

With these challenges in mind, we set out to design a platform that would enable: (i) live imaging a dynamic and asynchonised process using reporter systems; (ii) isolating individual events for downstream molecular and functional characterisation at specific timepoints. The platform had to fulfil two requirements: cells had to be cultured in contact with extracellular matrix (ECM), without attaching to the microfluidic device directly; and channels had to be controlled individually so that specific cells or cell aggregates of interest could be extracted independently.

Hydrogel microbeads provide an ideal system to keep cells in contact with ECM, without direct attachment to a dish or surface. Previously, it was shown that ES cells can self-renew and initiate the process of differentiation in agarose-only beads^5^. Here, we optimise conditions for encapsulating cells in 3D agarose-fibrin hydrogel beads. We also build a microfluidic platform that enabled us to retain in culture or extract for analysis individual microbeads by media flow, without the need of enzymatically dissociate the matrix. Finally, we show that this platform can be used to follow the asynchronous exit from the ES cell state over time, and determine the functional properties of cells at specific points during that process.

## RESULTS AND DISCUSSION

### 3D agarose-fibrin matrix mini-screen

Agarose is biologically inert and does not provide specific cell adhesion or ECM retention. Therefore, we sought to optimise the cell encapsulation platform by providing a biologically active scaffold component. Thrombin can polymerise fibrinogen molecules to form fibrin scaffolds that are highly biocompatible and able to bind various ECM proteins, including laminin^6,7^. Moreover, fibrin can be remodelled and digested by cells over time providing a system to potentially escape compression associated with cell division^7^, while it’s stability can be increased by covalent modification^8^. However, excess remodelling can eventually lead to disintegration of the scaffold. To strike a balance between biocompatibility, degradability and stability, we made use of agarose-fibrin composite matrices, where agarose provides the non-degradable scaffold.

To identify the optimal agarose/fibrin scaffold composition for ES cell differentiation in hydrogel beads, we designed a 3D matrix screen, for which we provide a template (Table S1). ES cells harbouring a Sox1::GFP (a neural marker) knock-in reporter were embedded in 96-well plates containing matrices of varying composition, and differentiated towards the neural lineage (FIG 1A). We varied the concentration of agarose, fibrinogen, thrombin and laminin in combination (TABLE 1) and examined the effect on differentiation efficiency. Some of the combinations led to scaffolds disintegration over time so that cells grew adherent at the bottom of the dish and were, therefore, poor candidates for stable encapsulation. In other conditions, however, the scaffold was stable and cells grew in 3D, embedded in the hydrogels. We assigned each well a matrix-integrity score of 1-4. Wells with a matrix integrity score of 4 contained only embedded aggregates, while matrix integrity scores of 1 indicated scaffolds that disintegrated. After 5 days, we dissolved scaffolds with agarase and dissociated cells with trypsin. The differentiation efficiency was analysed by flow cytometry (FIG 1A). All conditions resulted in Sox1 upregulation in >70% of cells (FIG 1B). The highest proliferation was observed when cells escaped encapsulation and grew preferentially adherent to the surface (matrix-integrity score of 1-2). Since removal of cells attached to a surface would necessitate enzymatic methods, we discarded such conditions. As expected, agarose concentration was the biggest determining factor for matrix integrity (FIG 1C). Conversely, in the gels with the highest matrix integrity, increased fibrinogen concentration showed a slight negative correlation with the percentage of Sox1::GFP positive cells and cell numbers (FIG 1D). Laminin and thrombin concentration did not show significant trends on their own (not shown). Taken together, our results indicate that the optimal conditions for ES cell differentiation were in hydrogel droplets formed under the higher agarose concentrations and lower fibrin. However, fibrin was still necessary, as omitting fibrin resulted in increased cell death after 3 days in culture (FIG 1E). Thus, we chose to carry forward the condition which resulted in the highest matrix stability score, cell proliferation and differentiation efficiency [1% Agarose, 0.075mg/ml Fibrinogen, 0.5U/ml Thrombin and 0.3μg/ml laminin, FIG 1B, red arrows].

**Table 1.** Concentration of components tested in combination on the matrix screen.

| Agarose<br>% | Fibrinogen<br>mg/ml | Thrombin<br>U/ml | Laminin<br>μg/ml |
| --- | --- | --- | --- |
| 1 | 0.5 | 0.25 | 0.6 |
| 0.85 | 0.2 | 0.5 | 0.3 |
| 0.70 | 0.075 | 1 |  |
| 0.50 | 0.025 |  |  |

**Figure 1.**
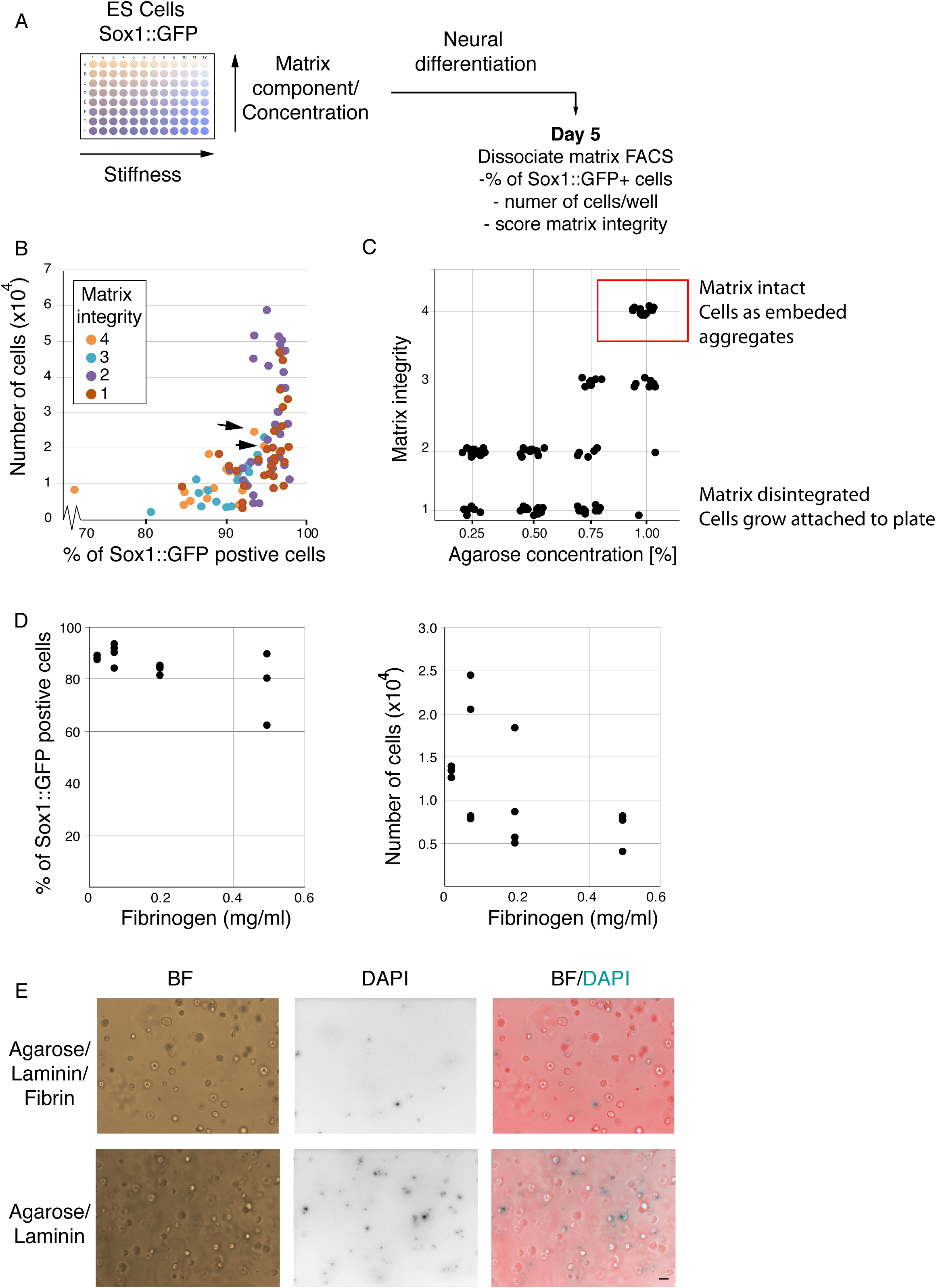
3D matrix mini-screen. A. Schematic of 3D matrix screen to determine hydrogel composition for efficient neural differentiation. B. Percentage Sox1::GFP positive cells and number of cells on day 5 of neural differentiation in droplets. Each dot represents a different matrix condition. C. Effect of agarose concentration on matrix integrity. D. Effect of fibrinogen concentration on percentage of Sox1::GFP cells and number of cells (E) after 5 days of differentiation. E. Cell survival in agarose/fibrinogen/laminin composite gels vs agarose/laminin-only matrices on day 3. DAPI is able to enter only dead or dying cells. Scale bar 100nm

### Functionalised microgels support multilineage differentiation of mES cells

Next, we examined whether we could generate hydrogel microspheres with the optimised matrix conditions. To this end, we used a flow-focusing microfluidic encapsulation device^5^. Low-melt agarose was dissolved and cooled to 37°C as previously described^5^, before adding fibrinogen and laminin. This solution was fed through one inlet of the encapsulation device. Newly formed agarose droplets were collected in tubes on ice. Basal media supplemented with Thrombin was added to the collection tube, and agarose-fibrin droplets were de-emulsified by addition of perfluorooctanol (PFO) (FIG 2A). Thrombin addition led to fibrin/fibrinogen incorporation and retention in the microgels (FIG 2B). The microspheres remained stable and did not attach to the dish.

**Figure 2.**
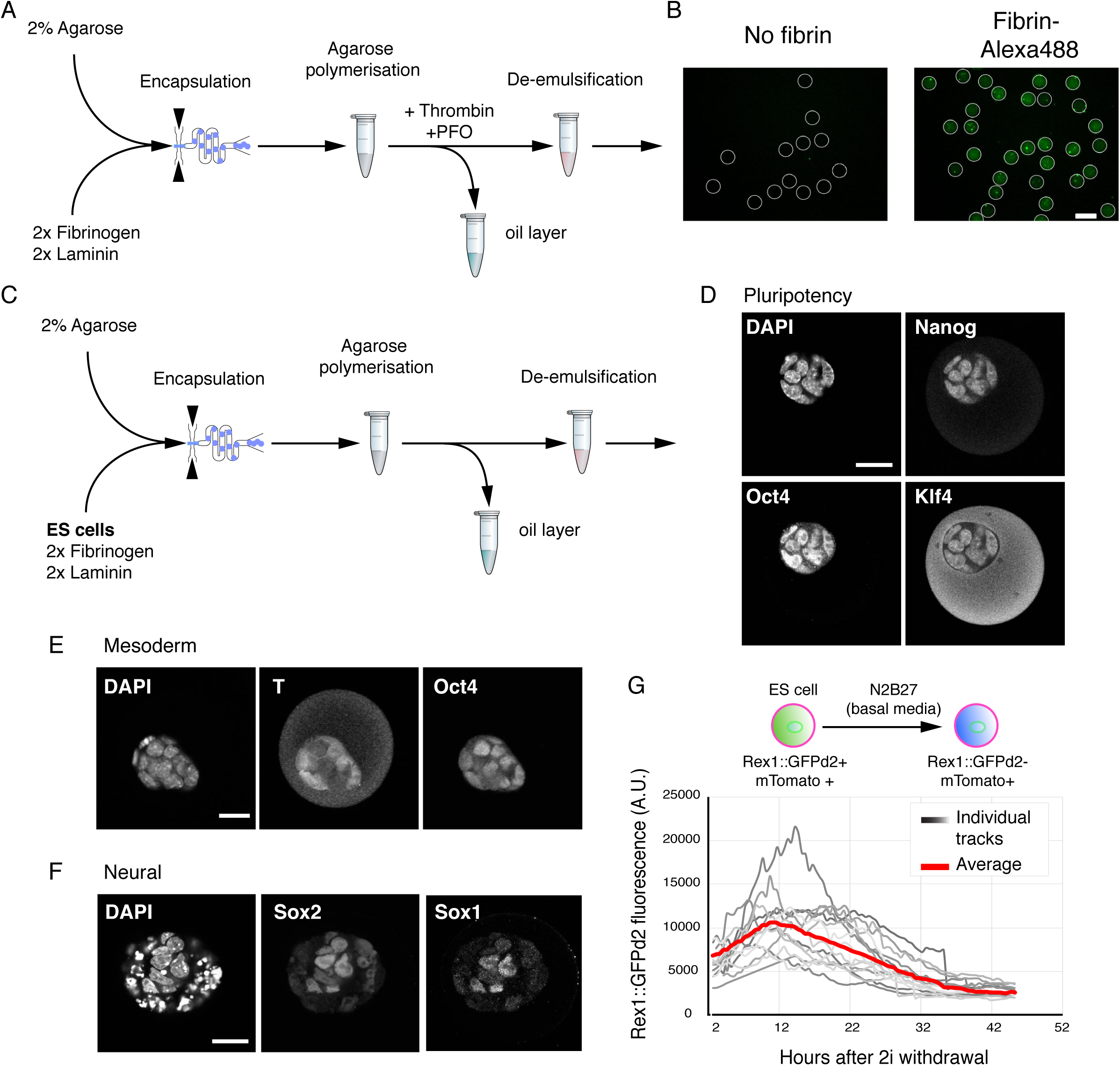
ES cell differentiation in functionalised hydrogel microspheres. A. Hydrogel droplet formation using a flow-focusing microfluidic devise. B. Fluorescently labelled fibrin is successfully retained in microgel beads. C. Strategy for encapsulating ES cells in hydrogel beads. D. Encapsulated ES cells maintained in ES cell media and fixed on 2 days for immunostaining show expression of pluripotency-associated transcription factors Nanog, Oct4 and Klf4. Nuclei were counterstained with DAPI. Scale bar 20μm. E. Encapsulated ES cells were placed in mesoderm-inducing media and fixed for immunostaining on day 3. Oct4/T double positive cells are early mesoderm progenitors, while Oct4+/T-cells are yet uncommitted cells. Scale bar 20μm. F. Encapsulated ES cells were directed towards the neural lineage and analysed on day 4 for the expression of transcription factors Sox1 and Sox2. Punctuated DAPI staining shows dead or dying cells. Scale bar 20μm. G. Live imaging of differentiating ES cells encapsulated in 3D hydrogels. Membrane Tomato (mTomato) expression was used to segment aggregates in 3D. The average aggregate level was determined for each droplet and plotted against the time after withdrawal of self-renewal signals (2i). Individual tracks and average are shown.

Next, we tested if the agarose-fibrin beads could sustain ES cell self-renewal and differentiation. 2% low-melt agarose solution was heated and subsequently cooled to 37°C. ES cells were resuspended in basal media (N2B27) supplemented with 2x fibrinogen and laminin solution. The agarose and cell suspension were mixed, and quickly passed through the cell encapsulation microfluidic device (FIG 2C). Encapsulated cells were de-emulsified as shown previously^5^, in a solution containing thrombin.

Under ES cell culture conditions, encapsulated cells proliferated at a rate similar to what previously reported in 3D^5^ and maintained expression of pluripotency markers Nanog, Klf4 and Oct4 (FIG 2D). When placed in mesoderm-inducing conditions, we could detect expression of the pan-primitive streak marker T(Bra) (FIG 2E), while in neural conditions we could detect Sox1 and Sox2 double positive neural precursor cells (FIG 2F). Finally, we tested whether the hydrogel droplets were amenable to live imaging. ES cells harbouring the ES cell-state reporter Rex1::GFPd2, and a constitutive membrane marker (mTomato), were encapsulated and allowed to exit the ES cell state in basal media (N2B27). 3D stacks of individual beads were acquired every 30 min and automatically segmented using the mTomato membrane marker to measure the total GFP level. This allowed us to track the downregulation of Rex1::GFPd2 as ES cells exited the naïve state (FIG 2G). In agreement with previous reports^4,9^, the downregulation of Rex1::GFPd2 was asynchronous, with cells in some gel beads downregulating the reporter before others.

Overall, the data demonstrate that agarose and fibrin can be successfully co-polymerised in hydrogel droplets, enabling support for both self-renewal and multi-lineage differentiation. Moreover, the system is compatible with live imaging. We noted that in some conditions, especially after 5-6 days of culture, cells towards the outside of the aggregate showed signs of cell death (e.g. fragmented nuclei), possibly due to compression. Since pluripotency transition experiments take place within 48 hrs of encapsulation, we do not anticipate this being a limitation. However, the matrix conditions could be further optimised for longer term experiments. Finally, the hydrogel droplets can easily be dissolved following agarase treatment (not shown). Since agarase is not cytotoxic, it can be added to culture media, without a need to deactivate or remove it for subsequent culture.

### Microfluidics for single droplet culture and retrieval

To achieve single droplet control we designed a 2-layer microfluidic platform using photolithography. A figure-8 channel layer contains individual droplet traps (FIG 3A, insert [i]), inlets and outlets. An auxiliary module contains an array of droplet traps (FIG 3A, insert [ii]) and multiple independent inlets/outlets for immunofluorescence. A second layer was superimposed onto the channel layer to control media flow by means of PDMS pneumatic (or Quake) valves^10^. To achieve complete channel closure upon valve actuation, the mould of the channel layer was constructed using a two-step process. First, non-compressible features were generated with SU8 2100. Then, features containing the channels under pneumatic control were superimposed onto the SU8 features, in AZ 40 XT and reflown by reheating to generate rounded profiles (FIG 3A, insert [iii] - ^11^).

**Figure 3.**
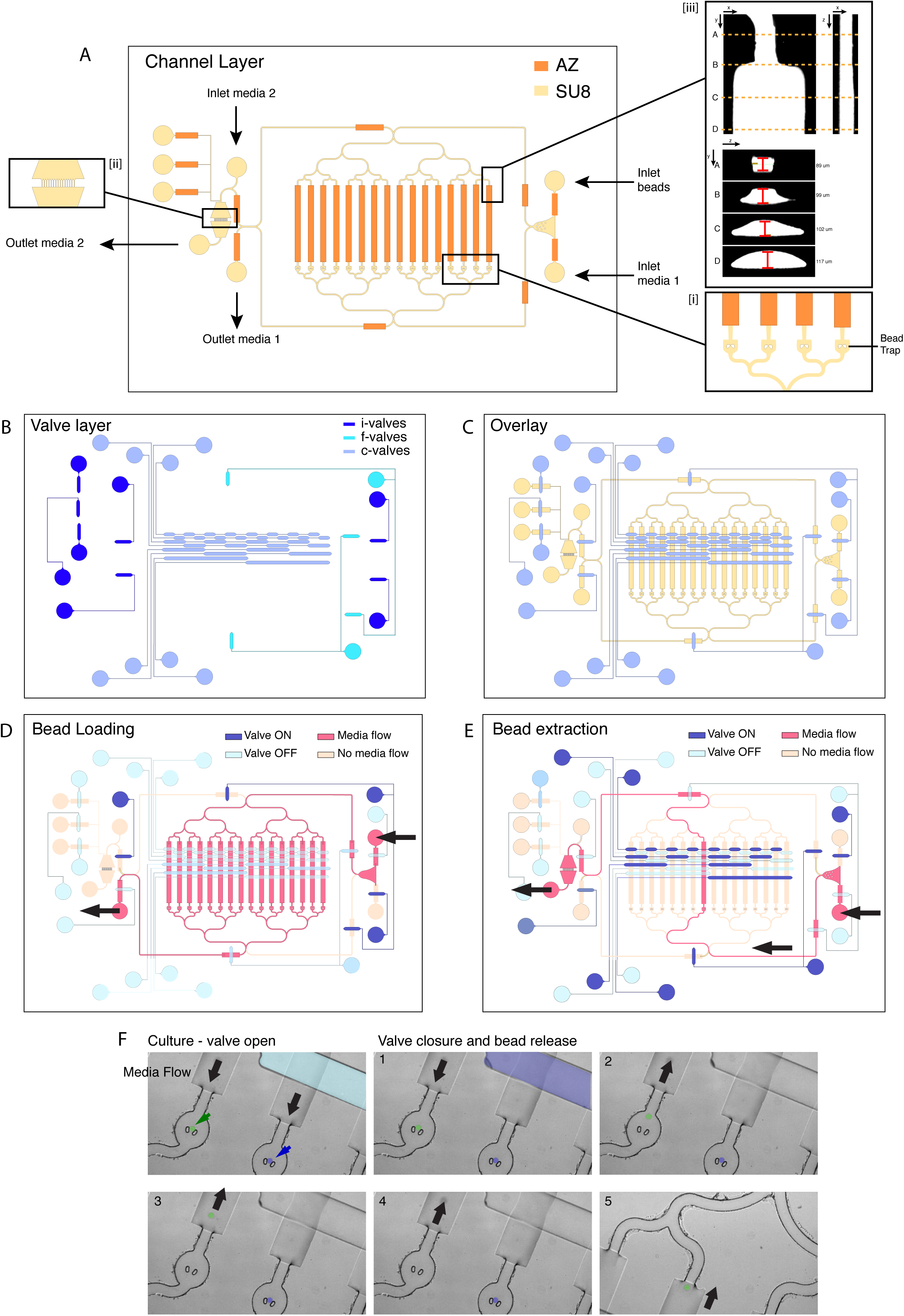
Microfluidic platform for independent capture and release of aggregates. A. Schematic of the channel (bottom) layer of the microfluidic platform. (i) shows individual bread traps, (ii) shows a magnified view of a secondary culture module, (iii) shows cross-section of the channels and height measurements. B. Schematic of the valve layers. C. Overlay of both PDMS layers. D. Valve actuation for loading beads. The media flows through all channels. E. Example combination of valves that result in removal of a single bead without affecting the remainder. F. Snapshots of live imaging showing valve actuation, and removal of a single bead (false colour in green), while a neighbouring bead (false colour blue) remains in position.

In the design, a number of valves control inlet/outlet access (FIG 3B, i-valves). These are actuated to prevent backflow and direct the media flow to specific outlets and/or auxiliary modules. Two valves control the direction of media flow into the trap module (FIG 3B, f-valves), while 8 valves provide single channel control (FIG 3B, c-valves). Activating different sets of valves enables controlling the direction of media flow during bead-loading and culture (FIG 3D), and removing individual beads (FIG 3E). Valve actuation is achieved through a pressure increase in the valve layer, which is controlled by a MUX microfluidic flow switch matrix (Evenflow), which can apply constant pressure over any combination of valves. For the droplet removal to be effective, the sequence and timing of valve actuations is controlled by a custom MATLAB graphical user interface (GUI). Within the GUI, each combination of valve actuations was mapped to a specific button relating to a corresponding bead trap. The user is able to dictate the flow direction, channel of interest and destination (outlet vs auxiliary module) through single mouse clicks on the GUI window.

We tested whether individual beads could be removed without displacing the remaining beads (FIG 3F). When the media flow is directed towards the trap (FIG 3F, panel 1) hydrogel beads are held in position. To extract an individual bead, the media flow was initially redirected towards the desired channel by closing off all the alternative channels (FIG 3F, panel 2). Next, the direction of media flow was reversed to extract the droplet of interest into a well of a 96-well plate (FIG 3F, panels 3-6). Notably, the neighbouring droplet remains in position.

Therefore, our platform allows for cell culturing under live imaging conditions of individual cells/beads, which can be independently removed. The platform, moreover, provides a very high degree of flexibility. The number of channels and droplets analysed can be doubled by adding two extra control valves (FIG 3B, c-valves). Moreover, the independent inlets can be used to carry out media change or drug administration at precise intervals of time.

### Live imaging of differentiation and single cell retrieval for functional studies

We designed an experiment to validate the platform. As cells exit the ES cell state, they lose the ability to self-renew/survive in ES cell maintenance media (2i/LIF). The ability to self-renew correlates with the expression of Rex1::GFPd2; cells that have extinguished Rex1::GFPd2 expression can no longer self-renew in selective ES cell culture conditions^4^. The phenotype of cells with intermediate levels of Rex1::GFPd2 is unknown. The analysis of these intermediates is complicated by the fact that differentiation is asynchronous (FIG 2G and^4^). To determine if cells with intermediate levels of Rex1::GFPd2 could self-renew or had irreversibly lost ES cell identity, we encapsulated Rex1::GFPd2/mTomato cells into functionalised gel bead. The self-renewal signals were withdrawn and cells maintained for 27 hrs, at which point they displayed a wide distribution of Rex1::GFPd2 expression levels^4^. We monitored Rex1::GFPd2 expression, before extracting individual cells/droplets from the platform (FIG 4A). Each gel bead was placed back in selective ES cell culture conditions (2i/LIF) and the capacity to form self-renewing colonies determined. We found that cells that retained detectable expression of Rex1::GFPd2, even if significantly lower than ES cell-controls, could still self-renew in 2i/LIF (FIG 4B, C). This indicates that the ability to self-renew in 2i/LIF conditions is lost only after cells have fully downregulated Rex1::GFPd2 expression. Therefore, our platform allows us to directly correlate reporter expression over time with the functional response to signalling pathways.

**Figure 4.**
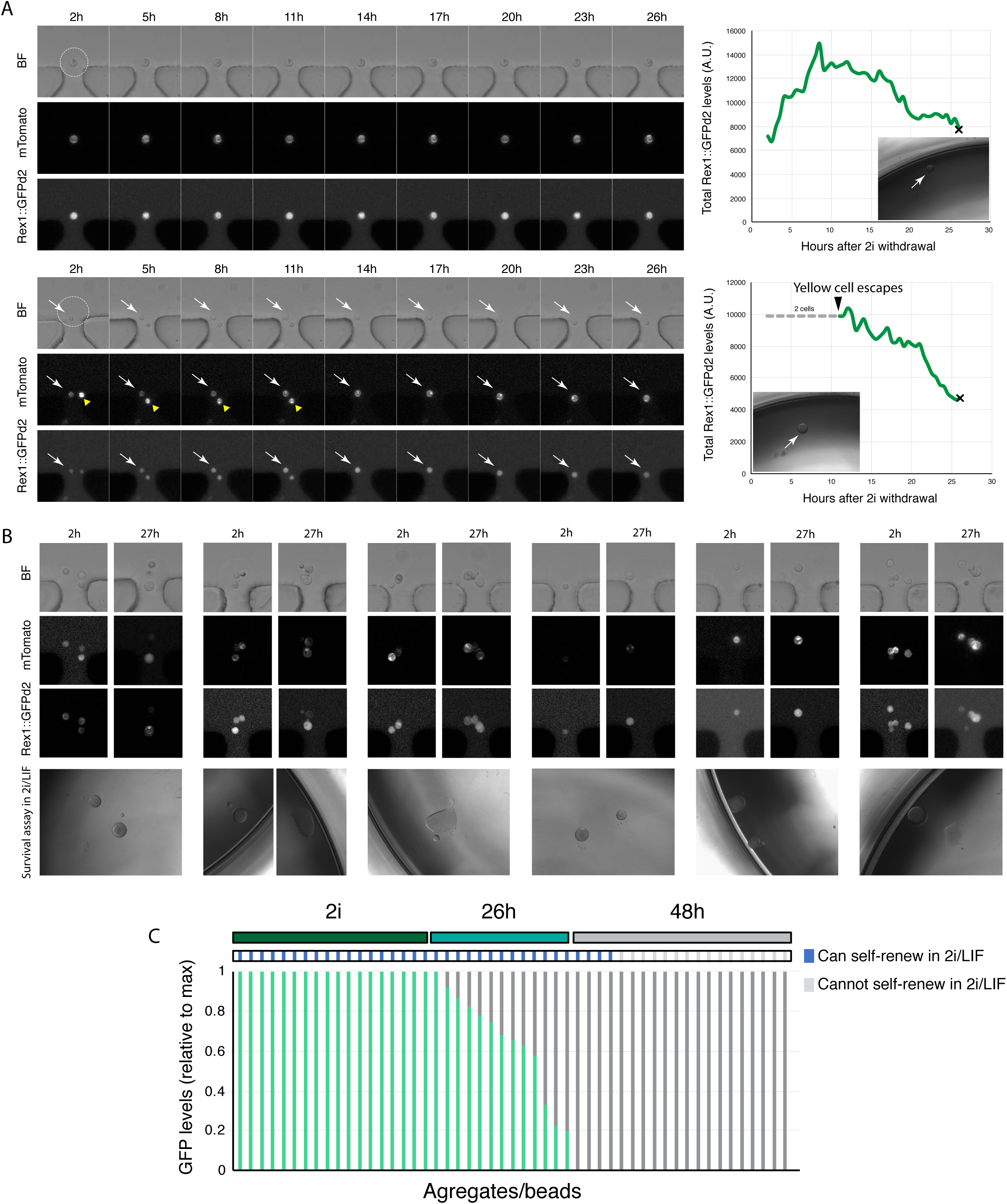
Combined platform for live imaging and retrieval of live cells for functional studies. A. Example tracks of single encapsulated cells, tracked over 27hrs during exit from the ES cell state. Cells were automatically segmented using mTomato and the total GFP level determined. Graphs on the right show total Rex1::GFP levels, normalised to cell/aggregate volume. ‘x’ denotes point of extraction and insert figure shows colony arising after 5 days in ES cell culture conditions (2i/LIF). B. Images of start and end point of live imaging of different hydrogel microbeads. C. Quantification of the total level of GFP, normalised to peak, expression, and the outcome of the self-renewal assay in 2i/LIF.

## CONCLUSIONS

In this study we present a versatile platform for integrating live imaging with individual cell retrieval. The platform offers the potential to link cell history with either molecular or functional characterisation.

We developed a hydrogel droplet encapsulation system that is highly customisable, such that the extracellular matrix composition, degradability and stiffness can be tuned. Under our optimised conditions, we could observe self-renewal and multilineage differentiation of mouse embryonic stem cells. The hydrogel matrix screen approach, for which we provide a template (TABLE S1), as well as the use of functionalised hydrogel droplets could both provide useful platforms for culture for different cell types.

Notably, the two-layer microfluidic platform allows capturing individual hydrogel droplets, culturing or differentiating them under living imaging conditions, and removing individual droplets at specific times. The design of the microfluidic platform, moreover, allows for doubling the number of captures hydrogel droplets for each two control valves that are added. Different reporter systems could be used to determine differentiation status, signalling pathway activity, cell cycle phase, or any other cellular feature, before cell extraction. The placement and removal of beads in individual traps is controlled by media flow and therefore does not require enzymatic treatments. The ability to isolate individual droplets while leaving the remainder unperturbed, could prove a useful system for isolating specific stages in processes that occur asynchronously or are of transient duration. Overall, we expect this platform to be adaptable to a wide range of applications for assessing dynamics of cellular processes.

## METHODS

### Mouse ES cell culture and differentiation

Mouse ES cell culture and differentiation was performed following previously described protocols ^9,12^. Briefly, cells carrying a constitutive tomato marker and a destabilised form of GFP under the control of the Rex1 promoter were routinely grown in 2i or 2iLIF [1μM PD0325901, 3μM CHIR99021, 100U/ml Leukaemia inhibitor factor (LIF)] in N2B27 basal media [1:2 DMEM/F12 (Signa-Aldrich D6421), 1:2 Neurobasal (Thermo Fisher 21103049), 0.5X B17 (Invitrogen 17504044), N2.BV (see below), 50μM b-mercaptoethanol (Thermo Fisher 31350), 2mM L-Glutamine (Thermo Fisher 25030081), 12.5μg/ml Insulin human recombinant zinc solution (Thermo Fisher 1258014)]. N2.BV was made in house [8.791mg/ml Apotransferrin (Sigma Aldrich T1147), 1X DMEM/F12, 0.66% BSA Fraction V, 3μM sodium Salenite (Signa Aldrich S5261), 1.688mg/ml Putrescine (Sigma Aldrich P5780), 2.08μg/ml Progesterone (Sigma Aldrich P8783)]. Cells were split every two days and plated at a density of 1.5 x 10^4^ cells/cm^2^. Differentiation in aggregates was carried out by placing hydrogel droplets in N2B27 (without 2i or LIF).

### Matrix screen

Low melt agarose (Lonza SeaPlaque 50105) was dissolved at twice the concentration required and kept at 37°C to prevent polymerisation. 2x Thrombin solution was added to each tube containing agarose. 50μl of agarose/thrombin were added to each well of a 96 well plate kept on a hot plate set to 37°C to prevent polymerisation. Next, basal media was supplemented with 2x laminin and 2x fibrinogen, and mixed with cells (final cell concentration per well 3500). 50μl of the media/laminin/fibrinogen and cell mixture was added to each well containing agarose/thrombin and mixed by pipetting while keeping the plate at 37°C. Once sufficiently mixed, the plate was placed at 4°C for 15min to allow agarose to polymerise. 100μl of N2B27 were added once the hydrogel was polymerised. Cells were differentiated for 5 days, before assigning each well a matrix integrity score (4=all cells in aggregates embedded in hydrogel, 3=most cells in aggregates embedded in hydrogel, 2=most cells growing adherent to the bottom of the well, 1=all cells growing adherent to the bottom of the well and no matrix visible). Next, media was carefully removed and replaced with 50ul of fresh N2B27 supplemented with Agarase (Thermo Fisher Scientific EO0461) and trypsin. Hydrogels were dissolved after 15minutes and gentle pipetting resulted in a single cell suspension. Differentiation efficiency was analysed by flow cytometry. A template for customising the matrix screen, alongside detailed instructions and calculations, is provided in TABLE S1.

### Immunofluorescence

Hydrogel beads were fixed in 4% PFA for 30min at room temperature, before washing twice with PBS for 5min each time. Aggregates were permeabilised and blocked for 2-4hrs in block solution (PBS, 0.3% Triton 100X (Sigma Aldrich T8787), 3% Donkey serum (Sigma Aldrich D9663). Primary antibodies were diluted in block solution and incubated overnight at 4°C under gentle rocking. Droplets were washed three times for 15min with PBST (PBS, 0.3% Triton 100X) before incubating with secondary antibodies diluted in block solution. After 3 washes of PBST for 15min each, beads were stored at 4°C in PBS to reduce background before mounting in VECTASHIELD antifade mounting media (Vector Laboratories H-1000). Images were acquired on a confocal laser scanning microscope (Leica SP5). The following primary antibodies were used: Nanog (1:200, eBioscience 14-5761-80), Oct4 C-10 (1:400, Santa Cruz sc5279), Klf4 (1:400, R&D AF3158), T(Bra) (1:300, R&D AF2085), Sox2 (1:200, eBioscience 14-9811-82), Sox1 (1:200, Cell Signalling 4194). Alexa Fluor-conjugated secondary antibodies were used (Life Technologies). Nuclei were stained with DAPI.

### Live imaging and analysis

Live imaging of agarose drops outside and inside the microfluid device was done using a Zeiss 710 confocal microscope using an 20x 0.5 EC Epiplan-Neofluar objective. A custom made-heated black box was used to surround the microscope and syringe pumps and avoid large temperature gradients. Ten z-stacks were acquired every 30min for GFP, mTomatoe and bright field. Image analysis was carried out in Imaris, using mTomatoe to automatically segment aggregates in 3D. At each timepoint, GFP levels were normalise to total aggregate volume.

### Hydrogel bead generation

Low melt agarose (Lonza SeaPlaque 50105) was dissolved at twice the required concentration (2%) in PBS by heating at 80°C, before reducing the temperature to 37°C. Cells were dissociated to obtain a single cell suspension at a concentration of 5-10 x10^6^ cells/ml in 2i supplemented with 0.15 mg/ml (2x) fibrinogen (Sigma Aldrich F4883) and 0.6μg/ml (2x) Laminin (Mouse purified, Merck Millipore CC095). The agarose solution was mixed 1:1 with the cells/fibrinogen/laminin in 2i solution and kept at 37°C briefly to avoid polymerisation. The encapsulation was carried out as previously described ^5^ with a few modifications. Briefly, chips was plasma bonded to glass coverslips and the inlets were immediately flushed with a solution of 1% PFOTS (Trichloro(1H,1H,2H,2H-perfluorooctyl)silane, Sigma Aldrich 448931) in HFE-7500 (3M Novec Engineered Fluid, Fluorochem 051243). For hydrogel droplet generation, one inlet was used for the agarose/cell suspension and one for the oil/carrier solution (HFE-7500 supplemented with 0.3% Pico-Surf 1 (Sphere Fluidics) which were flown through at 6μl/min and 30μl/min respectively. Newly formed hydrogel droplets were collected on ice and polymerised for 15min. Cell culture media (either 2i or N2B27 alone) was supplemented with 0.5U/ml Thrombin (Sigma Aldrich T4393) and added to the bead/oil emulsion. Deemulsification was carried out by adding 1H,1H,2H,2H-perfluorooctanol (PFO) (Alfa Aesar, B20156) directly to the oil layer, quickly vortexing and extracting the top aqueous layer containing the hydrogel droplets.

### Microfluidic platform generation

Devices are fabricated from two layers of PDMS. The base layer is formed by spin coating 20:1 (base:curing agent) PDMS (Slygard 184, Dow Corning) over the channel layer silicon wafer described above. This is sufficient to cover the SU8 (MicroChem, UK) and AZ (MicroChem, UK) features to create a thin PDMS membrane. We find that 450 RPM is sufficient enough to produce a robust membrane whilst being thin enough to deform as a valve. This layer is cured at 60°C for 6 hours.

Initially, a silicon wafer is spin coated with SU8 2100 at 3100 RPM to achieve a 90μm layer before 5 minute baking at 65°C and 30 minutes at 95°C. To create the ‘SU8’ features of FIG 3A, a photomask of similar design (ref JD) was used with a MJB-4 contact mask aligner (Karl Suss, Munich, Germany) to expose the SU8 coated wafer for 18s (365-405nm, 20mW cm^-2^). Following a subsequent post expose bake of 15 minutes at 95°C, the wafer is developed in propylene-glycol-mono-methyl-ether-acetate (PGMEA) (Merk Millipore) for 7 minutes. Residue is removed with isopropanol and the wafer is baked at 200°C to complete the SU8-layer fabrication.

The SU8 featured wafer was then spin coated with AZ 40 XT at 900 RPM. A slow-ramped bake from 65°C-126°C was performed over 20 minutes before the spin and bake process was repeated to achieve a double thickness layer. A photomask similar to the ‘AZ’ designs of Fig. A was used in the MJB-4 for exposure. The wafer is carefully aligned so the channels of the mask fit between the SU8 channels which are visible though the AZ layer. Exposure is two rounds of 90 seconds at 365-405nm, 20mW cm^-2^. The wafer is post-exposure baked at 105°C then immersed in AZ 726 developer (ref) with occasional water rinses until the features are revealed. To achieve the rounding of the valve sections, the wafer is then finally baked for 5 minutes at 110°C to reflow to a maximum height of 115-120μm (measured with stylus profilometer, Dektak, Bruker, MA, USA).

To produce a mould for the valve layer, SU8 2025 was spun across a silicon wafer at 1700 RPM to produce a feature height of 35μm. Bake and exposure protocols followed the manufacturer’s guidelines.

The valve layer is formed of 5:1 PDMS poured over the valve layer mould to a thickness of 1-1.5 cm and cured for 1 hour at 60°C. A scalpel is used to cut the PDMS chip around the outside of the features and free from the mould. A 1.5mm biopsy punch is then used to create holes (indicated by the circular sections of Fig. B) for fluidic access to the valve channels. The valve and channel layers are then exposed, feature side up, to oxygen plasma for 20s at 100W (Diener Electronic GmbH and Co. KG, Germany). Using a stereo microscope, the two layers are aligned as in Fig. C and brought into contact to bond together. Following a further 1-hour bake at 60°C, a scalpel is used to cut around the valve layer and through the channel layer to the device may be pealed from the wafer. Fluidic access is enabled through a 1.5mm biopsy punch at the circular sites of FIG. 3A. To complete the fabrication process, the PDMS device is bonded to a glass slide (76×52×1.2mm, Fischer Scientific) using the oxygen plasma protocol described above before baking at 60°C for one hour.

### Microfluidic platform set up and operation

To prepare microfluidic chips for use, the bonded PDMS and glass devices are submerged in sterile PBS and placed into a vacuum desiccator. The chips are held under vacuum over-night to replace air within the channels with PBS. Once filled, the chips are placed on the microscope stage for experimental set up. The stage is equipped with a heating plate set to 37°C for optimal cell culture temperature. Over the chip we are able to fix a transparent cover connected to a CO_2_ source which can be calibrated to supply (LEVELS). The combined effect of the heating and atmospheric control is to effectively replicate the conditions of a cell culture incubator.

For fluidic control, each of the valve inlets are connected via FEP tubing to the MUX multiplexing unit. For fluidic delivery, the channel inlets are connected via FEP tubing to a syringe pump (Nemesys, Cetoni). All equipment to control valves, flow and pressure are housed within a purpose-built cabinet containing a heater set to 37°C. This ensures media entering the chip does not undergo any temperature change between the syringe pump and the outlet. The chip is initially equilibrated with 1 ml of culture media via a constant flow of 5 ml/hr provided by the syringe pump.

## ACKNOWLEDGEMENTS

CM and ACH were supported by a Leverhulme Trust Research Grant (RPG-2016-418) and ERC grant (772798), TNK was supported by an AstraZeneca Graduate Studentship, CCA was supported by an MRC grant (MR/M011089/1), FH is a H2020 ERC Advanced Investigator [695669]. The Cambridge Stem Cell Institute receives core funding from the Wellcome Trust and the Medical Research Council. AS is a Medical Research Council Professor. KC was supported by a Royal Society University Research Fellowship.

## AUTHOR CONTRIBUTIONS

Conceptualisation, CM, ACH, AS and KC; Methodology, CM, ACH, CCA; Formal analysis, CM; Investigation, CM, ACH and TNK; Writing, CM, ACH, KC; Supervision, FH, AS, KC.

